# Rapid and Cost-Effective Preparation of a RuBisCO-Rich Protein Fraction from Dried Leafy Biomass

**DOI:** 10.64898/2026.06.05.730420

**Authors:** Alasdair DJ Freeman, Caroline A Evans, Kang Lan Tee, Tuck Seng Wong

**Affiliations:** School of Chemical, Materials and Biological Engineering, University of Sheffield, Sir Robert Hadfield Building, Mappin Street, Sheffield S1 3JD, United Kingdom; National Alternative Protein Innovation Centre (NAPIC), United Kingdom

**Author notes:** Address correspondence to: **Prof. Tuck Seng Wong**.

**Keywords:** RuBisCO, ribulose-1,5-bisphosphate carboxylase/oxygenase, alternative protein, plant protein, protein extraction

## Abstract

Ribulose-1,5-bisphosphate carboxylase/oxygenase (RuBisCO), the most abundant protein on Earth, is an attractive and sustainable food ingredient owing to its favourable nutritional and techno-functional properties. Leafy vegetables are particularly rich sources of RuBisCO; however, large-scale vegetable production generates substantial quantities of residual biomass throughout agri-food supply chains. Drying is widely used to stabilise this biomass and facilitate storage, transport, and handling, yet most reported RuBisCO extraction methods have been developed for fresh material and are poorly suited to dried feedstocks. Here, we present a simple, scalable, and cost-effective process for the recovery of food-grade RuBisCO from dried leafy biomass. Using spinach, rocket, and kale as model systems, efficient protein extraction was achieved from both freshly dried leaves and commercially available leaf powders without the need for resource-intensive processing. Application of the method to spinach yielded approximately 75 mg of high-purity RuBisCO per 100 g fresh-leaf equivalent, corresponding to an extraction efficiency of ∼70%, which increased to ∼90% following supplementation with 20 mM CaCl_2_. The recovered protein fraction also exhibited favourable foaming capacity and foam stability, demonstrating its potential as a functional food ingredient. This work provides a practical route for the valorisation of dried vegetable residues and supports the development of circular, waste-to-value supply chains for sustainable plant protein production.

## 1. INTRODUCTION

Ribulose-1,5-bisphosphate carboxylase/oxygenase (RuBisCO), the key enzyme of the Calvin-Benson cycle, catalyses the fixation of atmospheric carbon dioxide into organic carbon by converting ribulose-1,5-bisphosphate and CO_2_ into 3-phosphoglycerate during photosynthesis. Owing to its central role in primary productivity, RuBisCO is the most abundant protein on Earth, with an estimated global standing stock equivalent to approximately 5 kg per person (Phillips & Milo, 2009). Beyond its biological significance, RuBisCO is increasingly recognised as a promising and sustainable protein ingredient for food applications due to its favourable amino-acid composition and techno-functional properties, including solubility, emulsification, and foaming capacity (Di Stefano et al., 2018; Grácio et al., 2023; Pearce & Brunke, 2023).

At the same time, substantial quantities of RuBisCO-rich biomass are discarded across agri-food supply chains. Global vegetable production generates approximately 525 million tonnes of waste annually, corresponding to around 25% of total output (Akindele et al., 2025). As leafy green tissues are particularly rich in RuBisCO, this waste stream represents a largely untapped and low-cost source of high-quality plant protein. Valorising leafy vegetable residues for RuBisCO extraction could therefore simultaneously address food protein demand, improve resource efficiency, and reduce environmental burdens associated with food waste.

However, most reported RuBisCO extraction strategies rely on resource-intensive processing steps that are poorly suited to the large volumes, high moisture content, and rapid perishability of leafy vegetable waste streams. In practice, stabilisation steps such as drying are often required to enable storage, transport, and industrial-scale handling of this biomass. Yet existing extraction protocols have largely been developed for fresh material and are not optimised for dried leaves, limiting their practical applicability. To address this gap, we developed a simple, scalable, and low-cost method specifically tailored for the efficient recovery of food-grade RuBisCO from dried leafy biomass, thereby enabling the practical valorisation of this abundant but underutilised resource.

## 2. MATERIALS AND METHODS

### 2.1 Materials

All chemicals were purchased from Sigma-Aldrich (part of Merck), Thermo Fisher Scientific, or Santa Cruz Biotechnology unless otherwise stated.

### 2.2 Sources of leafy biomass

Baby spinach leaves were purchased from a local supermarket and dried overnight in an oven (Binder) at 50°C. The dried leaves were either processed immediately or stored at −20°C until further use. Powdered dried spinach leaves were purchased from Buy Wholefoods Online (BWFO) and stored at room temperature.

### 2.3 Preparation of RuBisCO-rich protein fraction

All procedures were performed at room temperature unless otherwise stated. Dried leaves (1 g) were resuspended in 20 mL water and incubated for 15 min. Insoluble plant material was removed by filtration through muslin cloth. The filtrate was clarified by centrifugation at 5,000 × g for 10 min. The pellet was discarded or, where indicated, re-extracted under identical conditions. Acid-washed activated charcoal (Sigma-Aldrich, C4386) was added to the supernatant at 2% (w/v) and mixed for 20 min to remove pigments and phenolic compounds. Charcoal was removed by centrifugation at 8,000 × g for 5 min, followed by filtration through grade 1 filter paper (Whatman). This treatment was repeated once when further clarification was required. Proteins were precipitated by adjusting the extract to pH 4.5 with HCl. The precipitated RuBisCO-rich fraction was collected by centrifugation at 12,000 × g for 15 min. The resulting pellet was freeze-dried and stored at −20°C until use.

To investigate the effect of pH on RuBisCO extraction, dried leaves were resuspended in buffer solutions instead of water. Extractions at pH 4.5 and 5.5 were performed using 25 mM citrate buffer; at pH 7.5, 25 mM Tris-HCl buffer; at pH 9.5, 25 mM glycine-NaOH buffer; and at pH 11.5, 25 mM sodium phosphate buffer.

To investigate the effect of divalent metal ions on RuBisCO extraction, CaCl_2_ (10–100 mM), CoCl_2_ (20 mM), MgCl_2_ (20 mM), MnCl_2_ (20 mM), NiCl_2_ (20 mM), MgSO_4_ (20 mM), or ZnSO_4_ (20 mM) were used.

Protein extracts were either analysed by sodium dodecyl sulfate-polyacrylamide gel electrophoresis (SDS-PAGE) for visualisation or quantified using the Pierce™ Bradford Plus Protein Assay (Thermo Fisher Scientific, 23236), as appropriate.

### 2.4 Total protein extraction from dried leaves

Total protein was extracted from dried leaves as described above, with the exception that the extraction buffer consisted of 100 mM Tris-HCl (pH 8.8), 1% (w/v) SDS, and 5 mM DTT. Dried leaves were resuspended in SDS-containing buffer and incubated at 95°C for 2 min, followed by solid-liquid separation by centrifugation. Where necessary, the pellet was subjected to a second extraction using the same buffer.

### 2.5 Denaturing polyacrylamide gel electrophoresis

Proteins in the RuBisCO-rich fraction were separated by molecular mass using SDS-PAGE. Gels were cast and run using a Mini-PROTEAN vertical electrophoresis system. Typically, 15 µL of protein extract was mixed with 5 µL of 4× loading buffer and heated at 94°C for 15 min prior to loading. Samples were resolved on a discontinuous gel comprising a 5% stacking gel (0.125 M Tris-HCl, pH 6.8) and a 15% resolving gel (0.375 M Tris-HCl, pH 8.8). Electrophoresis was performed at 200 V for 50 min in Tris-glycine-SDS running buffer. Following separation, gels were fixed in 10% (v/v) acetic acid and 20% (v/v) methanol, rinsed thoroughly with water, and stained with 0.1% (w/v) Coomassie Brilliant Blue R-250 in 50% methanol and 10% acetic acid. Gels were subsequently destained using acetic acid/methanol solution until clear background was achieved. Molecular weight standards (PageRuler Broad Range Protein Ladder, Thermo Fisher Scientific; 5–250 kDa) were used for size estimation. Gel bands were quantified using Fiji software (Schindelin et al., 2012).

### 2.6 Native polyacrylamide gel electrophoresis

Protein complexes were separated under non-denaturing conditions using native polyacrylamide gel electrophoresis (native PAGE). Samples were resolved on a 5% resolving gel prepared in 0.375 M Tris-HCl (pH 8.8). Typically, 6 µL of sample was mixed with 2 µL of native loading buffer (lacking SDS and reducing agents) and loaded without heat denaturation. Electrophoresis was carried out at 120 V for 1 h in Tris-glycine running buffer (25 mM Tris, 192 mM glycine). Following separation, gels were fixed in 10% (v/v) acetic acid and 20% (v/v) methanol for 15 min and stained with Coomassie Brilliant Blue R-250. Gels were subsequently destained using the same fixing solution until a clear background was obtained. Protein sizes were estimated using a NativeMark™ unstained protein standard (Invitrogen), covering a molecular mass range of 20–1236 kDa.

### 2.7 Absorbance measurement

Polyphenols and chlorophyll in the extracts were monitored by UV-Vis absorbance spectrophotometry using a Jenway 7615 spectrophotometer. Spectra were recorded from 300–700 nm using 1 mL samples in 1-cm pathlength cuvettes. Data were analysed using OriginPro software.

### 2.8 Sample preparation for protein identification

The RuBisCO-enriched fraction was resolved by SDS-PAGE and visualized using Coomassie Brilliant Blue staining. The gel band corresponding to the RuBisCO large subunit was excised and subjected to proteolytic digestion with trypsin to generate peptides for mass spectrometry analysis (Shevchenko et al., 2006). Peptide sequences were subsequently identified using nano-flow liquid chromatography coupled to tandem mass spectrometry (LC–MS/MS).

### 2.9 Peptide sequence analysis using LC–MS/MS

Peptides were separated on an Easy-Spray C18 column (75 µm × 50 cm) using a two-step gradient from 94% solvent A (0.1% formic acid in water) to 50% solvent B (0.1% formic acid in 80% acetonitrile) over 30 min at 300 nL/min, with the LC system (U3000 RSLCnano, Thermo Scientific) coupled to a hybrid quadrupole-orbitrap mass spectrometer (Q Exactive HF, Thermo Scientific). The mass spectrometer was programmed for data-dependent acquisition with 10 product ion scans (resolution 30,000, scan range 200–2000 m/z, automatic gain control (AGC) 1e5, maximum injection time 60 ms, isolation window 2 Th, fixed first mass 100, normalised collision energy 27) per full MS scan (resolution 120,000, scan range 375–1500 m/z, intensity threshold 3.3e4, minimum AGC 2e3, maximum injection time 60 ms) with 20 s dynamic exclusion time. Mass spectrometry data were processed using the peptide-mapping workflow with the Host Cell Protein option in BioPharma Finder™ 5.2 (Thermo Fisher Scientific). Protein identification was performed against the *Spinacia oleracea* proteome (UniProt Proteome ID: UP001155700; 29,244 entries; downloaded 09/08/2025). Trypsin was specified as the cleavage enzyme, with static modification of cysteine carbamidomethylation and N-terminal acetylation, and variable oxidation of methionine and tryptophan. Peptide assignments required both MS and MS/MS data, a mass tolerance of 5 ppm, and a minimum confidence score of 95%. Assignments corresponding to metal adducts, nonspecific protease activity, gas-phase-generated ions, or unspecified modifications were excluded (Millán-Martín et al., 2023).

### 2.10 Foaming capacity and foaming stability evaluation

RuBisCO was extracted as described above at a solid-to-liquid ratio of 1 g dried leaves per 10 mL extraction buffer. For foamability measurements, 15 mL of protein extract was homogenised for 5 min at 13,000 rpm using a T 18 digital ULTRA-TURRAX® (IKA) equipped with an S18 N - 10 G dispersing tool (IKA). The resulting foam was immediately transferred to a 50 mL graduated cylinder, and the foam volume was recorded to determine foaming capacity (FC) according to the equation below:

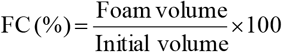

Foam stability (FS) was assessed by measuring the foam volume again after 30 min, using the equation below:

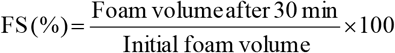

All experiments were performed in triplicate.

## 3. RESULTS & DISCUSSION

### 3.1 A simple and cost-effective method for preparing a RuBisCO-rich protein fraction

In developing a method for RuBisCO extraction from leafy biomass, several criteria were prioritised: (i) compatibility with the intended downstream applications of the protein, (ii) procedural simplicity to enable scalability and broad adoption, (iii) cost-effectiveness to support economic feasibility at larger scales, and (iv) safety, particularly for potential food-related uses. These considerations guided our selection of process steps and reagents.

Our extraction workflow comprises four stages: (i) leaf drying at 50°C, (ii) aqueous extraction of RuBisCO using water, (iii) removal of chlorophyll and phenolic compounds using acid-washed activated charcoal, and (iv) acid precipitation of the target protein fraction (Figure 1A). Compared with previously reported methods (Table 1), this approach is both time- and cost-efficient and does not require specialised equipment or expensive reagents.

**Table 1.**
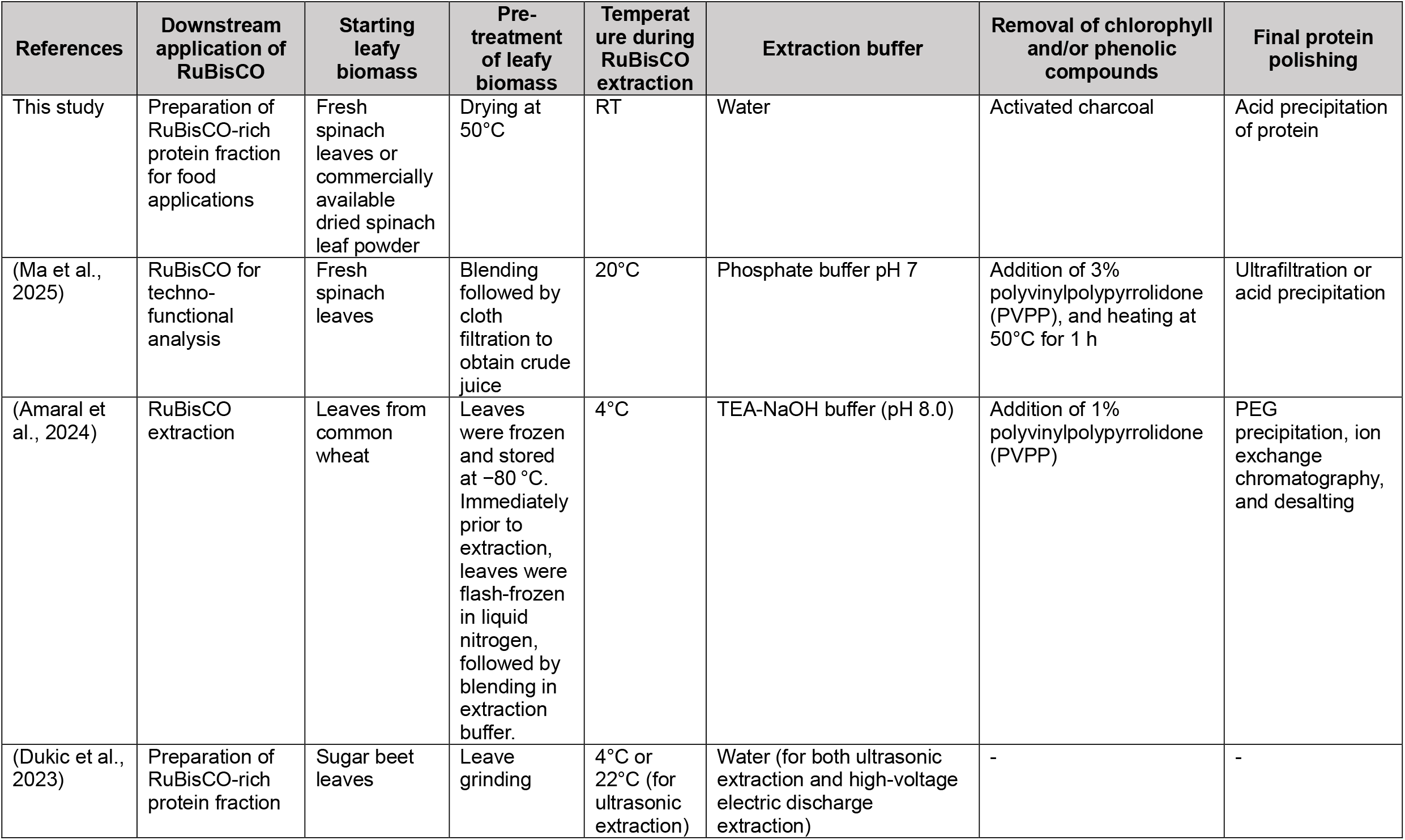
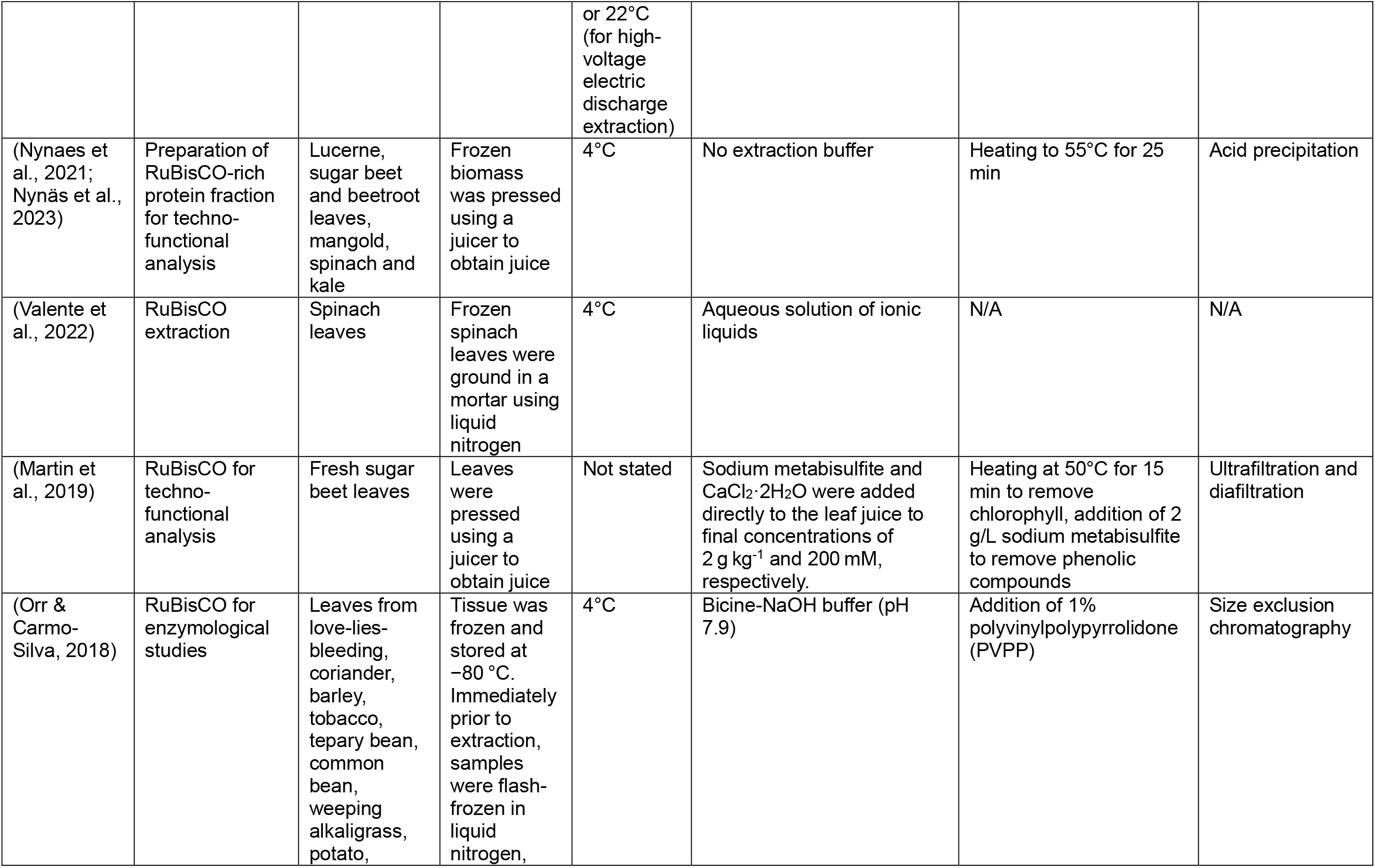
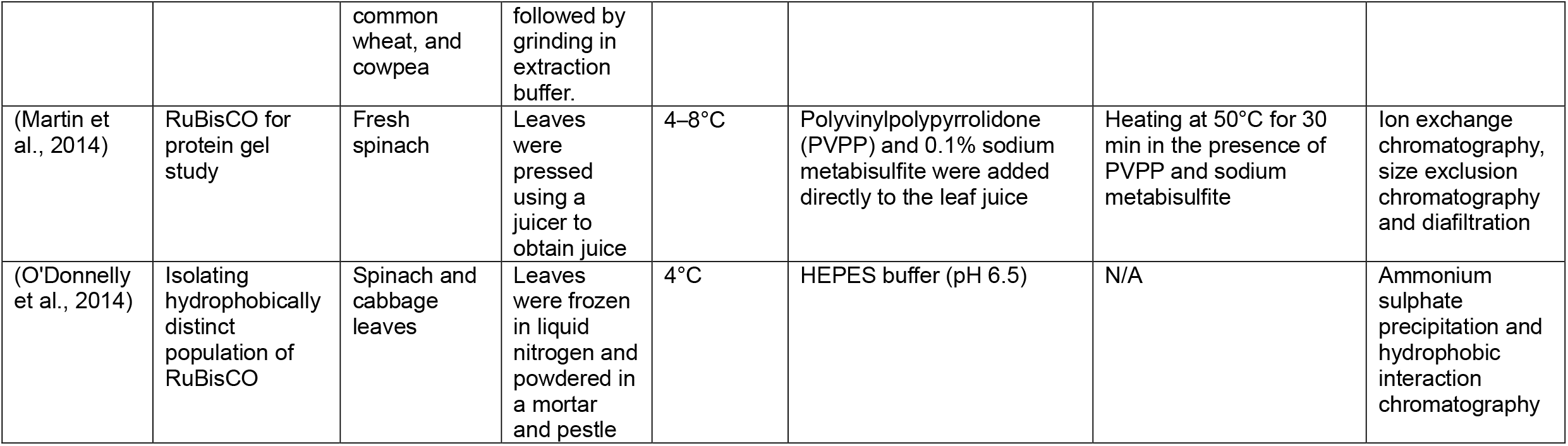
Comparison of RuBisCO extraction protocols reported in the literature and the method developed in this study.

**Figure 1:**
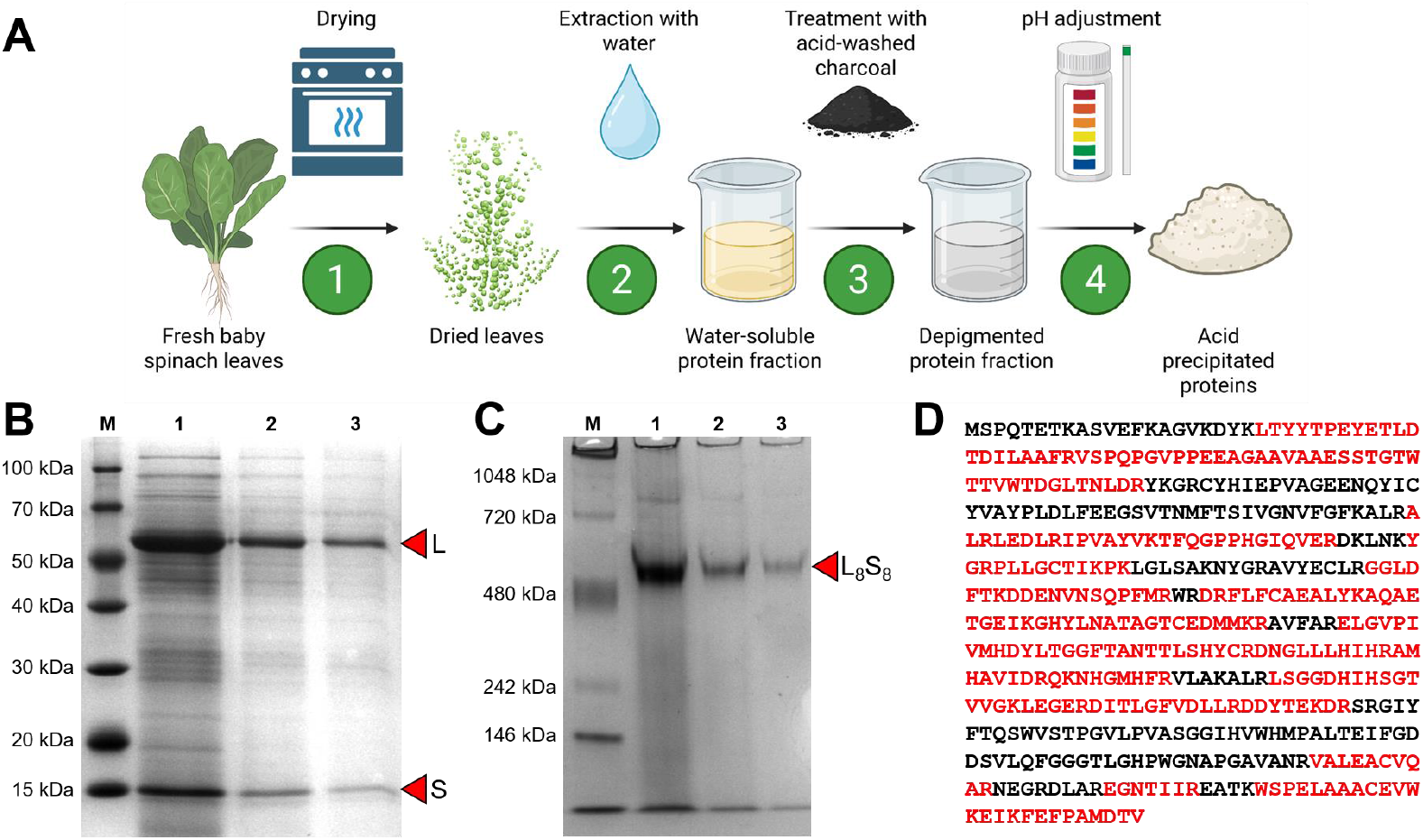
**(A)** A simple, cost-effective, and scalable process for preparing a RuBisCO-rich protein fraction, comprising four key steps. Step 1: Fresh baby spinach leaves were dried overnight at 50°C; this step can be omitted when commercially available dried leaf powder is used. Step 2: The dried leaf material was rehydrated with water to extract water-soluble proteins. Step 3: The extract was treated with acid-washed activated charcoal to remove chlorophyll, phenolic compounds, and other pigments. Step 4: The pH was adjusted to 4.5 to precipitate RuBisCO. This final precipitation step may be omitted when a soluble protein fraction is required for downstream applications. **(B)** SDS-PAGE analysis of proteins extracted from dried leaves by sequential water extractions: first extraction (lane 1), second extraction (lane 2), and third extraction (lane 3). The large and small subunits of RuBisCO are indicated as ‘L’ and ‘S’, respectively. Most RuBisCO was recovered during the first extraction. **(C)** Native PAGE analysis of proteins from the same sequential extractions (lanes 1–3). The RuBisCO holoenzyme complex is indicated as ‘L_8_S_8_’. **(D)** The protein band corresponding to the RuBisCO large subunit (panel B, lane 1) was excised and subjected to identification by LC–MS/MS. Database searching against the *Spinacia oleracea* proteome (UniProt Proteome ID: UP001155700; 29,244 entries) confirmed the identity of the large subunit of RuBisCO. Peptides identified by MS are highlighted in red within the full-length amino acid sequence of the protein.

### 3.2 RuBisCO extraction from baby spinach leaves

In C_3_ plants grown under low-stress conditions, including spinach (*Spinacia oleracea*), RuBisCO accounts for approximately 20–30% of total leaf-soluble protein (Galmés et al., 2014). Its abundance is highest during the leaf expansion phase, when photosynthetic capacity and protein synthesis are most active (Kudo et al., 2020). Accordingly, baby spinach leaves were selected as a model system for method development. As spinach is also widely used in related studies (Table 1), this choice enables direct comparison with previously reported data.

Drying leaves at 50°C disrupted cellular integrity, facilitating RuBisCO release without the need for mechanical or chemical disruption (Figure 1A). Removal of structural water during drying induces tissue deformation and compromises cell wall and membrane integrity, thereby enhancing protein extractability (Lewicki & Pawlak, 2003). In addition to aiding extraction, drying is a simple and widely used preservation strategy for plant materials, including medicinal plants (Nakra et al., 2025), making it well suited to scalable processing. Unlike previous studies that employed drying at 60°C (Rawiwan et al., 2023), the temperature of 50°C was selected in this study to remain below the reported melting temperatures (T_m_) of RuBisCO (Table 2), which are typically ≥60°C, thereby minimising the risk of thermal denaturation. Overnight drying at 50°C consistently resulted in a 90–95% reduction in biomass due to water loss, yielding a more compact and manageable material for subsequent handling and processing.

**Table 2.**
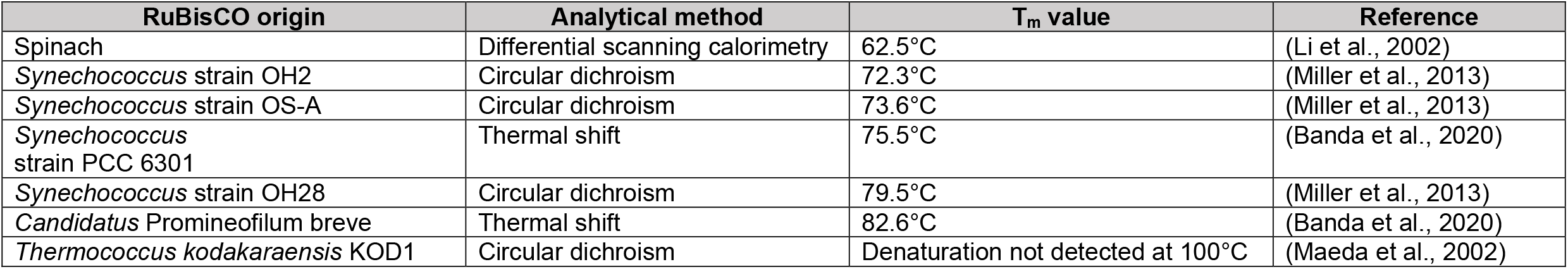
Reported melting temperatures (T_m_) of RuBisCO from various organisms.

| RuBisCO origin | Analytical method | $T_m$ value | Reference |
| --- | --- | --- | --- |
| Spinach | Differential scanning calorimetry | 62.5°C | (Li et al., 2002) |
| *Synechococcus* strain OH2 | Circular dichroism | 72.3°C | (Miller et al., 2013) |
| *Synechococcus* strain OS-A | Circular dichroism | 73.6°C | (Miller et al., 2013) |
| *Synechococcus* strain PCC 6301 | Thermal shift | 75.5°C | (Banda et al., 2020) |
| *Synechococcus* strain OH28 | Circular dichroism | 79.5°C | (Miller et al., 2013) |
| *Candidatus Promineofilum breve* | Thermal shift | 82.6°C | (Banda et al., 2020) |
| *Thermococcus kodakaraensis* KOD1 | Circular dichroism | Denaturation not detected at 100°C | (Maeda et al., 2002) |

Rehydration of dried spinach leaves with water efficiently extracted water-soluble proteins (Figure 1B). According to the Osborne classification (Osborne, 1924), this fraction corresponds to the albumin fraction and was highly enriched in RuBisCO complex, with a pH of 6.8. Both the large subunit (UniProt: P00875; calculated M_r_ ≈ 53 kDa; theoretical pI 6.13) and the small subunit (UniProt: P00870; calculated M_r_ ≈ 14 kDa; theoretical pI 5.41) were readily detected (Figure 1B). Native PAGE analysis confirmed that RuBisCO predominantly remained in its native L_8_S_8_ holoenzyme configuration, with an apparent molecular mass of approximately 550 kDa (Figure 1C), indicating preservation of structural integrity during the drying process. Proteomic analysis further confirmed the identity of the large subunit (Figure 1D), achieving 61% peptide sequence coverage.

Although aqueous extraction was effective, complete liquid recovery was not achieved, with approximately 15–20% of the extract retained within the leaf matrix. Sequential extractions with fresh water increased overall RuBisCO recovery (Figures 1B and 1C).

To estimate RuBisCO extraction efficiency, aqueous extraction was benchmarked against extraction with 1% (w/v) SDS, which was assumed to achieve complete protein recovery (Supplementary Figure S1A). Densitometric analysis of the RuBisCO gel bands indicated that a two-step aqueous extraction achieved an efficiency of approximately 70%. Bradford assay measurements showed that ∼75 mg of total soluble protein was recovered from 100 g of fresh leaves.

We next investigated whether RuBisCO recovery could be further enhanced by NaCl extraction, which targets the globulin fraction according to the Osborne classification. NaCl extraction was performed either as a secondary extraction following aqueous extraction or as the primary extraction step (Supplementary Figure S1B). In both cases, NaCl did not improve RuBisCO recovery, regardless of the salt concentration used (data not shown). Instead, NaCl extraction increased the level of co-extracted protein impurities, likely due to the solubilisation of additional salt-soluble proteins. These results demonstrate that NaCl supplementation is unnecessary for efficient RuBisCO extraction and may compromise product purity.

### 3.3 RuBisCO extraction from commercial dried leaf powder

Our extraction method was readily applicable to commercially available dried leaf powder. SDS-PAGE analysis revealed a protein profile comparable to that obtained from freshly prepared dried leaves (Figure 2A). Interestingly, RuBisCO extracted from the commercial material largely retained its native holoenzyme configuration, as confirmed by native PAGE (Figure 2B), despite storage at room temperature for more than six months. Structural stability of the complex was further evaluated by urea denaturation. Incubation in up to 2 M urea for 10 min preserved the intact holoenzyme, whereas partial dissociation was observed at 3 M urea, and complete loss of the complex occurred at 4 M urea (Figure 2C). Together, these findings indicate that leaf drying provides an effective strategy for the long-term preservation of RuBisCO in plant biomass. To our knowledge, the retention of native RuBisCO complex integrity following prolonged ambient storage of dried leaf material has not been previously reported.

**Figure 2:**
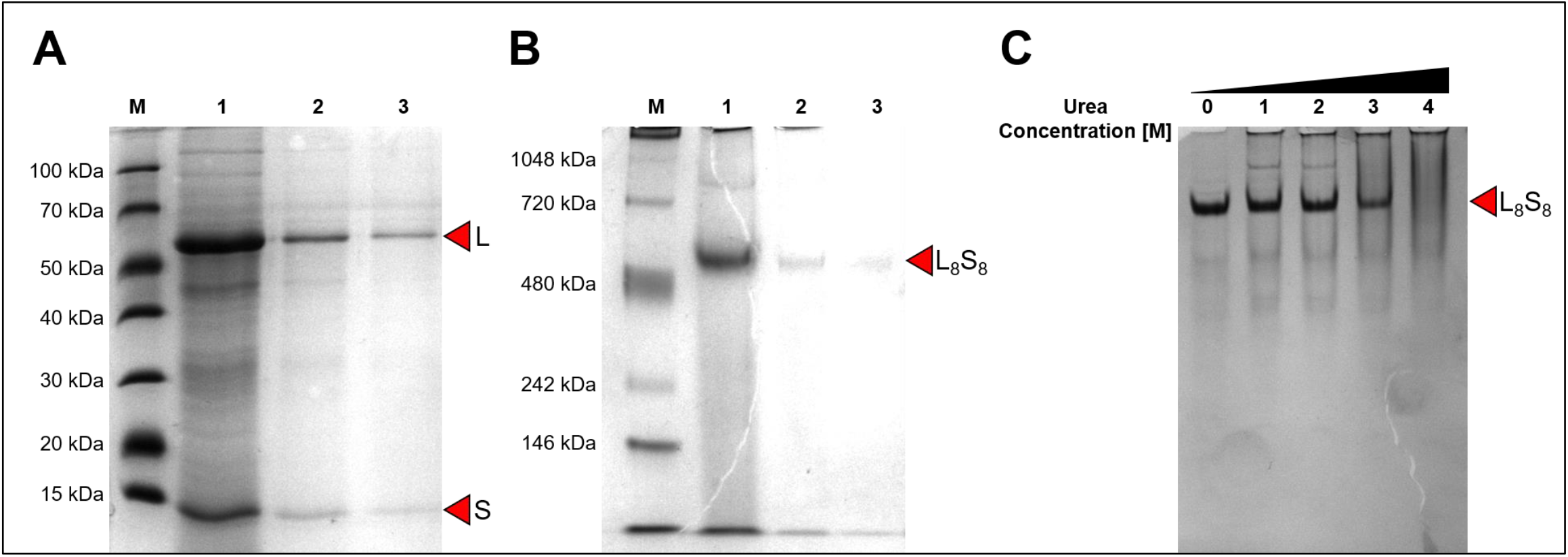
**(A)** SDS-PAGE analysis of proteins extracted from commercially available pre-dried spinach leaf powder by sequential water extractions: first extraction (lane 1), second extraction (lane 2), and third extraction (lane 3). The large and small RuBisCO subunits are indicated as ‘L’ and ‘S’, respectively. Most RuBisCO was recovered during the first extraction. **(B)** Native PAGE analysis of proteins from the same sequential extractions (lanes 1–3), showing the RuBisCO holoenzyme complex, indicated as ‘L_8_S_8_’. **(C)** The stability of the RuBisCO holoenzyme was assessed by incubating the protein extract (panel B, lane 1) with increasing urea concentrations from 0 mM (lane 1) to 4 M (lane 4). The complex began to dissociate at 3 M urea and was no longer detectable at 4 M.

### 3.4 RuBisCO extraction from other leaf types

In addition to spinach, the same extraction protocol was successfully applied to rocket leaves (*Eruca sativa*) and kale (*Brassica oleracea*) (supplementary Figure S2), yielding comparable protein profiles and demonstrating that the method is broadly applicable and not restricted to a specific plant type.

### 3.5 Pigment reduction

Aqueous extraction of soluble proteins using water produced a yellowish liquid (Figure 3A), which may be visually unappealing to consumers. UV-Vis spectroscopic analysis showed strong absorbance in the 300–450 nm region and weaker absorbance between 650 and 700 nm, consistent with the presence of chlorophyll and other plant-derived pigments (Figure 3B). Chlorophyll characteristically absorbs light in the blue–violet (400–450 nm; maxima at 430 nm for chlorophyll *a* and 453 nm for chlorophyll *b*) and red (640–670 nm; maxima at 662 nm for chlorophyll *a* and 642 nm for chlorophyll *b*) regions of the visible spectrum (McConnell et al., 2010). In addition to contributing to colour, chlorophyll is associated with earthy or grassy off-flavours that may reduce consumer acceptability (van Lelyveld & Smith, 1989). Therefore, to improve both visual appearance and sensory quality, activated charcoal and bentonite were evaluated as potential adsorbents for chlorophyll removal from the protein extract.

**Figure 3:**
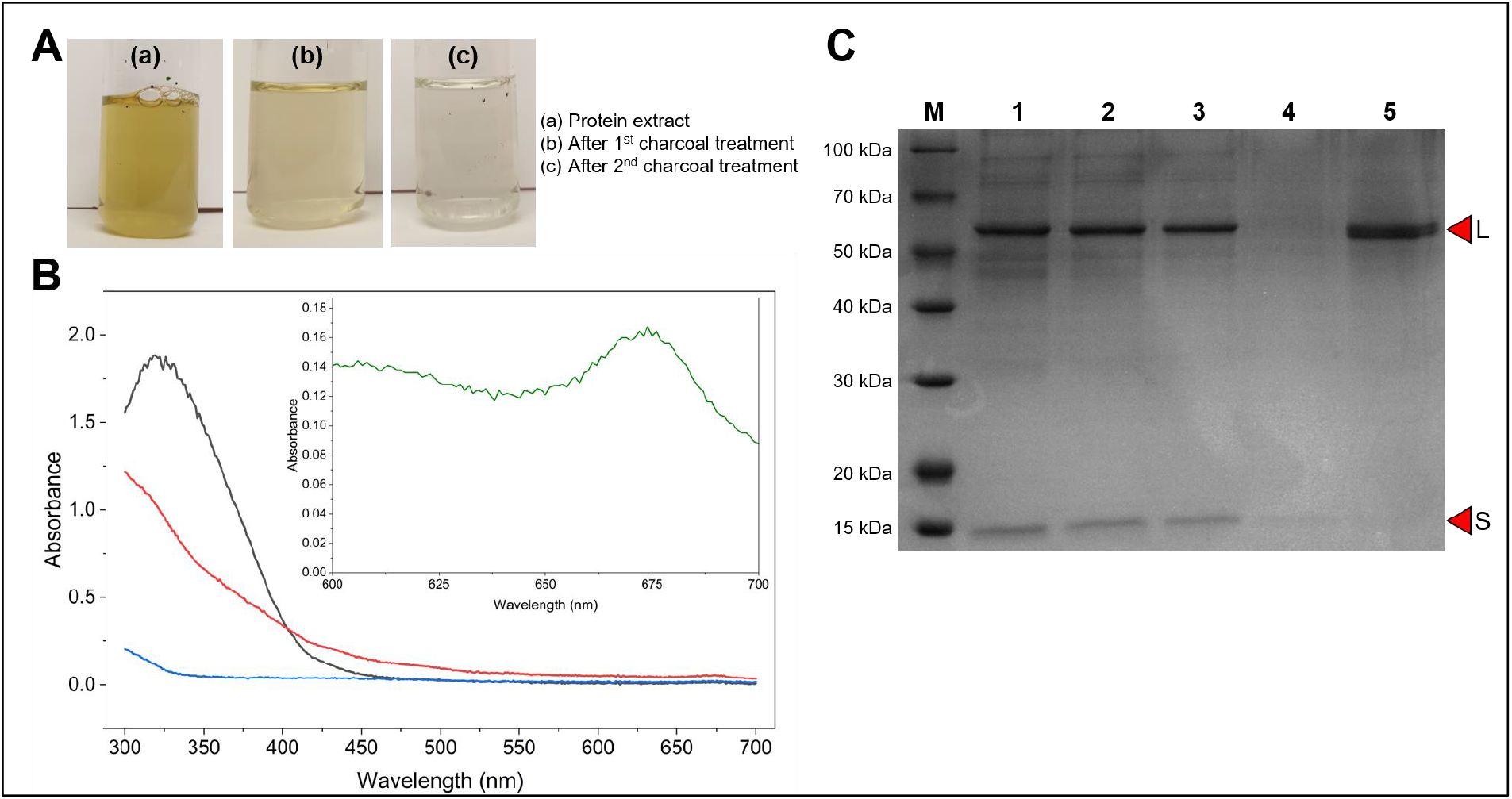
**(A)** The protein extract from dried leaves exhibited a yellowish colour (a). Following the first treatment with acid-washed activated charcoal, the colour intensity decreased markedly (b). After a second sequential charcoal treatment, only a faint yellow colour remained (c), indicating substantial removal of chlorophyll and other pigments. **(B)** UV-Vis spectra (300–700 nm) of a 10-fold diluted untreated protein extract (black), the undiluted extract after the first charcoal treatment (red), and the undiluted extract after the second charcoal treatment (blue). The inset shows the undiluted untreated extract (green). Charcoal treatment markedly reduced absorbance in the 300–400 nm and 650–700 nm regions, consistent with removal of chlorophyll and related pigments. Treatment efficiency was quantified using the absorbance at 326 nm, corresponding to the maximum of the untreated sample. **(C)** SDS-PAGE analysis of the untreated extract (lane 1), after the first charcoal treatment (lane 2), after the second charcoal treatment (lane 3), after pH adjustment to 4.5 to precipitate RuBisCO (lane 4), and after resolubilisation of the precipitated protein at pH 12 (lane 5). Charcoal treatment did not result in detectable protein loss, as indicated by comparable band intensities in lanes 1–3. Acid precipitation effectively enriched RuBisCO, as shown in lanes 4 and 5.

Following two rounds of charcoal treatment, a substantial reduction in plant pigments was observed, with approximately 95% removal after the first treatment and 99.5% removal after the second, as calculated from absorbance at 326 nm (Figures 3A and 3B). Importantly, charcoal treatment did not result in any detectable protein loss (Figure 3C). After clarification, RuBisCO was precipitated by adjusting the extract pH from approximately 6.8 to 4.5. Freeze-drying of the precipitate yielded 46 mg of protein per 100 g of fresh leaves.

Although bentonite has been reported as an effective adsorbent for chlorophyll removal (Tang et al., 2023), it proved unsuitable here because RuBisCO exhibited strong binding to the matrix, leading to near-complete depletion of the protein from the extract (supplementary Figure S3).

### 3.6 Effect of pH on RuBisCO extraction

We next examined the influence of extraction pH on RuBisCO recovery by replacing water with buffers spanning a broad pH range. At pH 4.5, no detectable RuBisCO was recovered (Figure 4A), consistent with our earlier observation that RuBisCO precipitates near its isoelectric point. In contrast, efficient extraction was achieved at pH 5.5 and above (Figure 4A). Native PAGE analysis confirmed that the extracted protein predominantly assembled into the native L_8_S_8_ holoenzyme complex across this pH range (Figure 4B).

**Figure 4:**
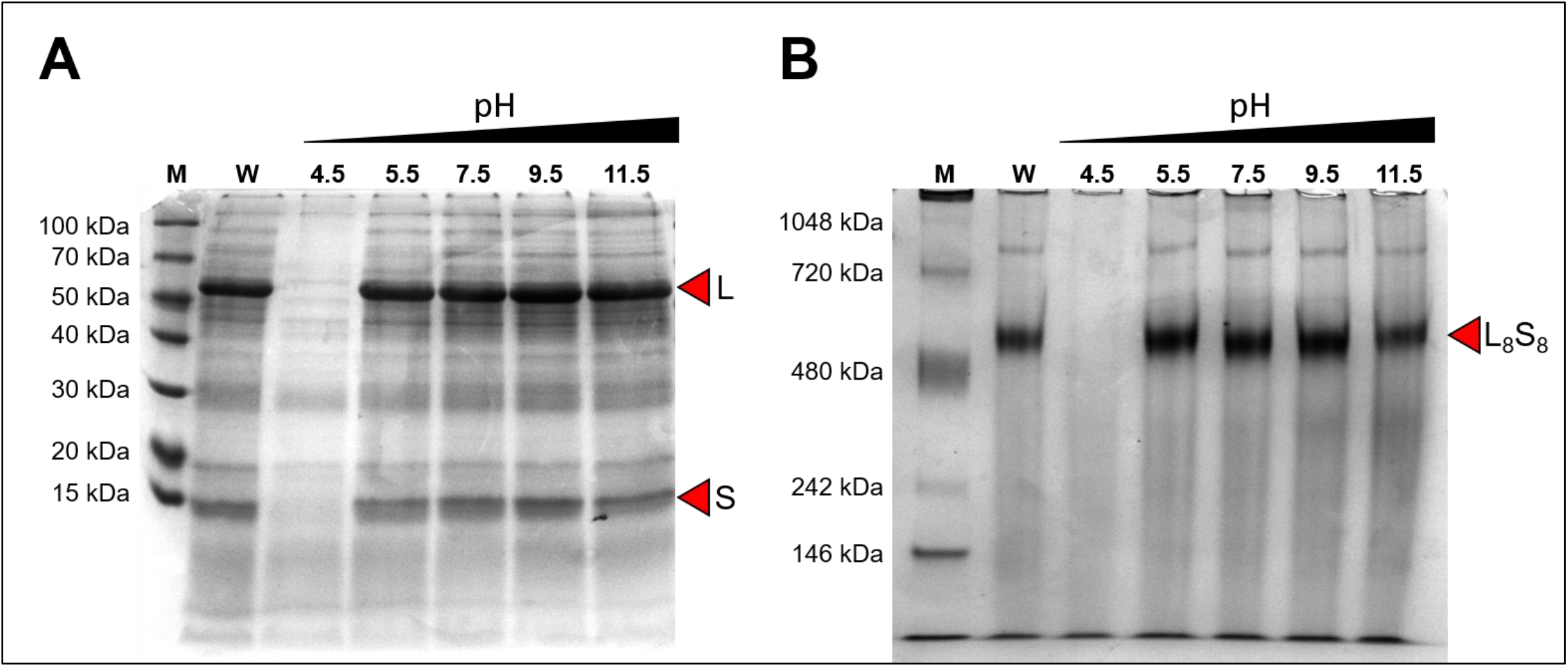
**(A)** SDS-PAGE analysis of proteins extracted from dried leaves using water (W) or buffers at pH 4.5, 5.5, 7.5, 9.5, and 11.5. With the exception of pH 4.5, all conditions effectively extracted RuBisCO. The large and small RuBisCO subunits are indicated as ‘L’ and ‘S’, respectively. **(B)** Native PAGE analysis of the same samples shown in panel A. The RuBisCO holoenzyme complex is indicated as ‘L_8_S_8_’. Reduced complex formation was observed at pH 11.5, suggesting partial disruption of the holoenzyme under highly alkaline conditions.

However, at pH 11.5, a noticeable reduction in intact complex formation was observed (Figure 4B), suggesting partial destabilisation under strongly alkaline conditions. This effect is likely due to high pH-induced chemical modifications of amino acids, including the formation of covalent cross-links such as lysinoalanine (LAL) and lanthionine (LAN) (Friedman, 1999), as well as racemization of serine from the L-to D-form (Friedman et al., 1984). Such modifications can compromise holoenzyme stability and explain the reduced assembly observed at very high pH. This observation also highlights a potential limitation of strongly alkaline extraction conditions, which are commonly employed in plant protein isolation (Hadinoto et al., 2024; Yao et al., 2023).

### 3.7 The effect of divalent metal ions on RuBisCO extraction

Motivated by proteomics studies demonstrating that Ca^2+^/phytate systems can selectively deplete RuBisCO from plant extracts (Krishnan & Natarajan, 2009; Sultan et al., 2013), we investigated whether supplementation of the extraction buffer with CaCl_2_ could improve RuBisCO recovery. CaCl_2_ is a generally recognised food additive that is widely used across a variety of food products, including baked goods, potato-based snacks, dairy products, beverages, juices, coffee, tea, condiments, jams, and processed meats.

To assess its effect, CaCl_2_ was added at concentrations ranging from 10 to 100 mM (Supplementary Figure S4A). Supplementation with 10 or 20 mM CaCl_2_ resulted in a substantial increase in RuBisCO recovery, reaching approximately 90% at 20 mM. This concentration corresponds to approximately 0.22% (w/v) CaCl_2_ and is below the practical upper limit of 0.4% (w/v) (≈36 mM) commonly considered acceptable in food-related applications (Bailone et al., 2022). In contrast, further increases in CaCl_2_ concentration led to a decline in RuBisCO recovery. This reduction is likely attributable to a salting-out effect (Liu et al., 2023), whereby Ca^2+^ ions shield charged residues on the protein surface, reducing electrostatic repulsion and promoting protein aggregation and precipitation.

To determine whether this effect was specific to Ca^2+^, we also evaluated other divalent metal ions, including Co^2+^, Mg^2+^, Mn^2+^, Ni^2+^, and Zn^2+^. Among these, Mn^2+^ produced a comparable enhancement in RuBisCO recovery (Supplementary Figure S4B), suggesting that the observed improvement is dependent on the identity of the divalent cation and may reflect ion-specific effects on protein solubility or extractability.

In addition to increasing RuBisCO recovery, extraction in the presence of 20 mM CaCl_2_ reduced absorbance across the 300–700 nm wavelength range (Supplementary Figure S4C), indicating lower co-extraction of plant pigments and other chromophoric compounds. This resulted in improved extract clarity.

### 3.8 Foamability of RuBisCO-rich protein solution

We investigated and compared the foaming properties of water-extracted, NaCl-extracted, and CaCl_2_-extracted RuBisCO-rich protein solutions (Table 3). All three samples exhibited good foaming capacity (FC) and foaming stability (FS), indicating that RuBisCO-rich fractions possess inherent surface-active properties that are retained across extraction conditions. No statistically significant differences in FC were observed among the three protein solutions.

**Table 3.** Foaming capacity (FC, %) and foaming stability (FS, %) after 30 min for RuBisCO-rich protein solutions prepared using different extraction buffers (water, 500 mM NaCl, and 20 mM CaCl_2_).

|  | **Water** | **500 mM NaCl** | **20 mM CaCl<sub>2</sub>** |
| --- | --- | --- | --- |
| Protein concentration (mg/mL), quantified using a Bradford assay | 0.76 | 0.70 | 0.84 |
| Foaming capacity (%), n = 3 | 171 ± 4 | 160 ± 12 | 182 ± 17 |
| Foaming stability (%) after 30 min, n = 3 | 45 ± 5 | 87 ± 0 | 53 ± 3 |

In contrast, FS was significantly higher in the NaCl-extracted sample. This enhancement may be attributed to salt-dependent modulation of protein-protein and protein-interface interactions, including changes in electrostatic screening, solubility, and interfacial film rheology, which together can enhance foam stability. Similar salt-induced effects on foam stability have been reported for other food proteins, including egg white (Yu et al., 2023) and dairy proteins (Zhang et al., 2024). In addition, the NaCl extraction may have co-extracted additional NaCl-soluble proteins, which could also have contributed to the observed changes in foaming behaviour.

We did not compare the measured FC and FS values with those reported for other RuBisCO preparations, as both parameters are highly sensitive to processing conditions, including the type of homogenisation equipment and the specific operating parameters used. Consequently, differences in instrumentation and processing settings can significantly affect foam formation and stability, limiting the validity of direct cross-study comparisons.

## 4. CONCLUSION

This study presents a simple, scalable, and broadly applicable method for the extraction of RuBisCO from dried leafy biomass. Water-based RuBisCO extraction offers the additional advantage of avoiding the introduction of extraneous compounds from the extraction buffer that could impart undesirable tastes or flavours. Furthermore, the absence of chemical extraction reagents eliminates the risk of introducing potentially toxic compounds into the final product, thereby enhancing food safety and preserving the suitability of the extracted RuBisCO for food formulation applications. As this work was established using defined model systems comprising unmixed plant materials, future work will evaluate the performance and robustness of the process when applied to heterogeneous and mixed leafy residues that more closely represent real-world waste streams. In addition, further characterisation of the nutritional quality and techno-functional properties of proteins extracted from mixed biomass, including solubility, emulsification, foaming, and gelation behaviour, will be essential to determine their suitability for diverse food applications.

## Abbreviations

BWFO: Buy Wholefoods Online
LAN: lanthionine
LAL: lysinoalanine
PAGE: polyacrylamide gel electrophoresis
PVPP: polyvinylpolypyrrolidone
RT: room temperature
RuBisCO: ribulose-1,5-bisphosphate carboxylase/oxygenase
SDS: sodium dodecyl sulfate

## ACKNOWLEDGEMENT

This study is funded by the UK National Alternative Protein Innovation Centre (NAPIC), which is an Innovation and Knowledge Centre funded by the Biotechnology and Biological Sciences Research Council (BBSRC) and Innovate UK (Grant Ref: BB/Z516119/1).

