## Supplementary information for "Rapid and Cost-Effective Preparation of a RuBisCO-Rich Protein Fraction from Dried Leafy Biomass"

**Figure S1: (A)** SDS-PAGE analysis of proteins extracted from freshly dried spinach leaves using either water or a buffer containing 1% (w/v) SDS as the extraction solvent. In both cases, protein extraction was performed in two sequential steps, denoted as the $1^{st}$ and $2^{nd}$ extracts. Extraction with 1% (w/v) SDS resulted in the co-extraction of a substantial amount of protein impurities compared with water extraction. **(B)** SDS-PAGE analysis of proteins extracted sequentially with water ($1^{st}$ and $2^{nd}$ extracts), followed by extraction with 0.5 M NaCl ($3^{rd}$ and $4^{th}$ extracts). This extraction strategy was compared with direct extraction using 0.5 M NaCl, performed in two sequential steps ($1^{st}$ and $2^{nd}$ extracts). The large and small RuBisCO subunits are indicated as 'L' and 'S', respectively.

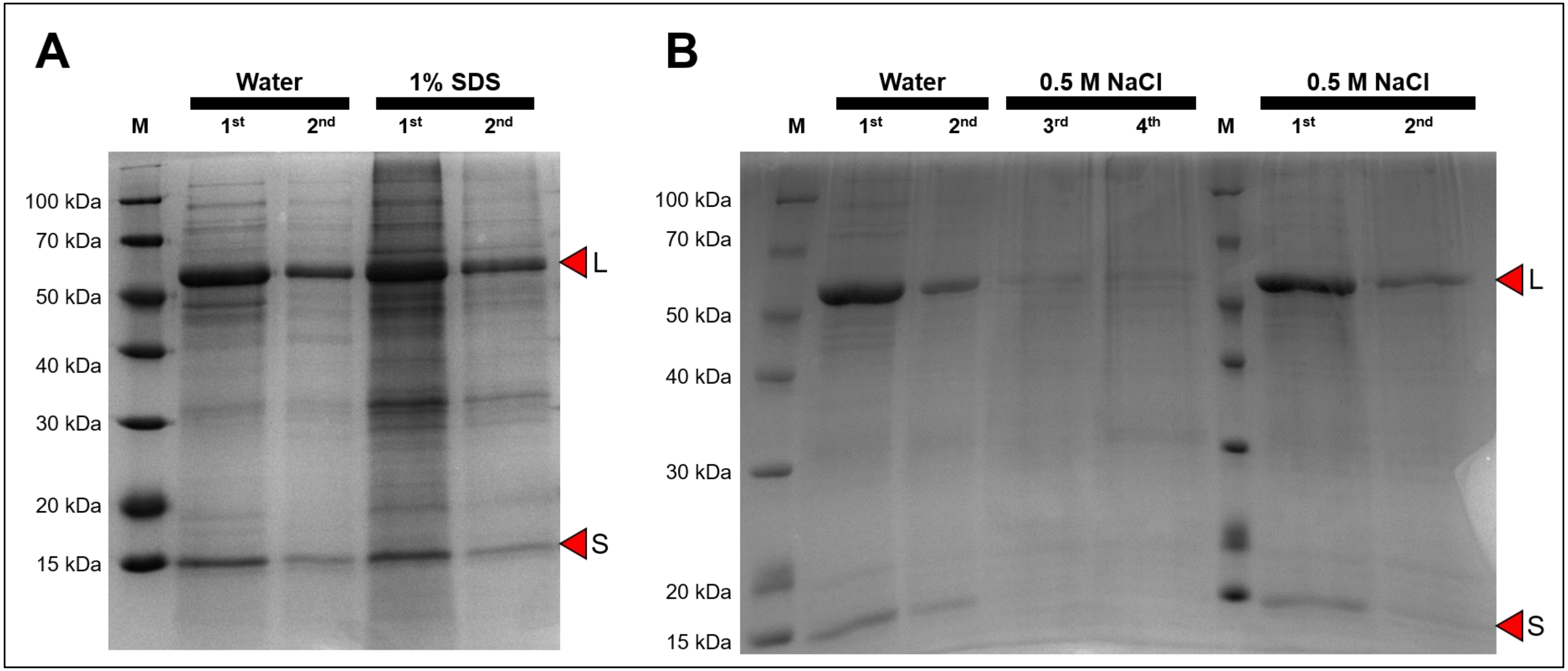

**Figure S2:** SDS-PAGE analysis of proteins extracted from rocket leaves **(A)** and kale leaves **(B)**. The large and small RuBisCO subunits are indicated as 'L' and 'S', respectively.

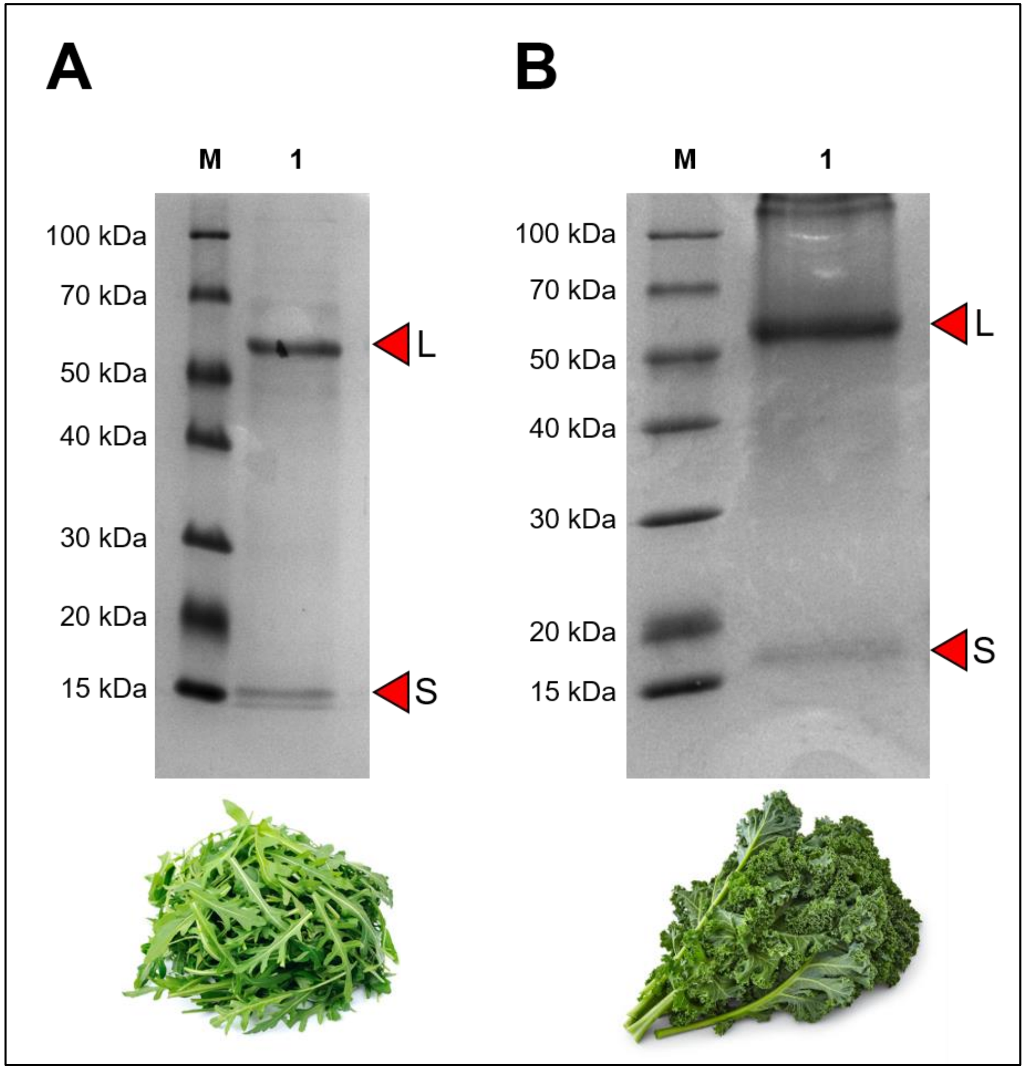

**Figure S3:** SDS-PAGE analysis comparing untreated protein extract (lanes 1–2) with bentonite-treated extract (lane 3). The large and small RuBisCO subunits are indicated as ‘L’ and ‘S’, respectively. Bentonite treatment resulted in a marked reduction in RuBisCO band intensity, indicating significant protein loss.

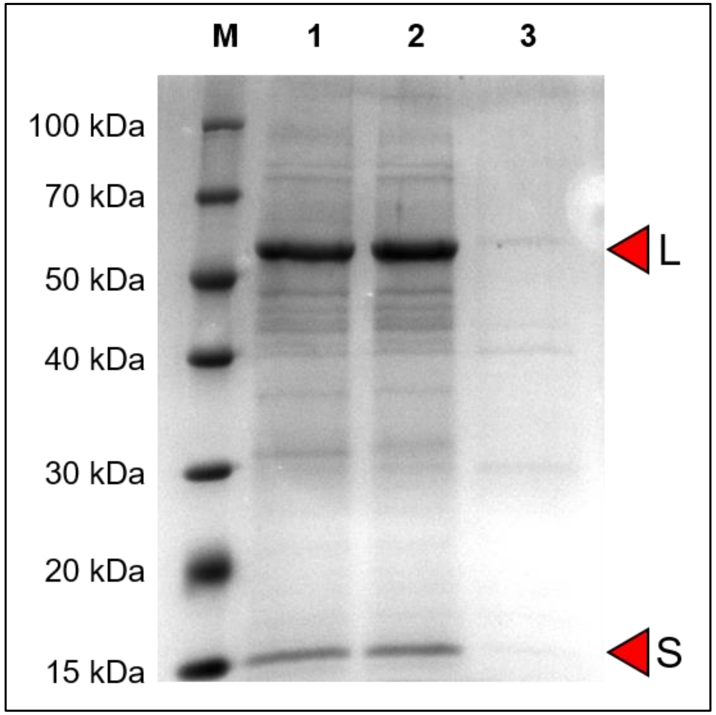

**Figure S4: (A)** SDS-PAGE analysis of proteins extracted from freshly dried spinach leaves using water (lane marked "W") or increasing concentrations of $CaCl_2$ (10–100 mM). **(B)** SDS-PAGE analysis of proteins extracted with water (lane marked "W") or with 20 mM solutions of divalent metal salts, including $CaCl_2$, $CoCl_2$, $MgCl_2$, $MnCl_2$, $NiCl_2$, $MgSO_4$, and $ZnSO_4$. The large and small subunits of RuBisCO are indicated as "L" and "S," respectively. **(C)** UV-Vis absorption spectra (300–700 nm) of protein extracts diluted 20-fold in water, comparing samples extracted using water or 20 mM $CaCl_2$.

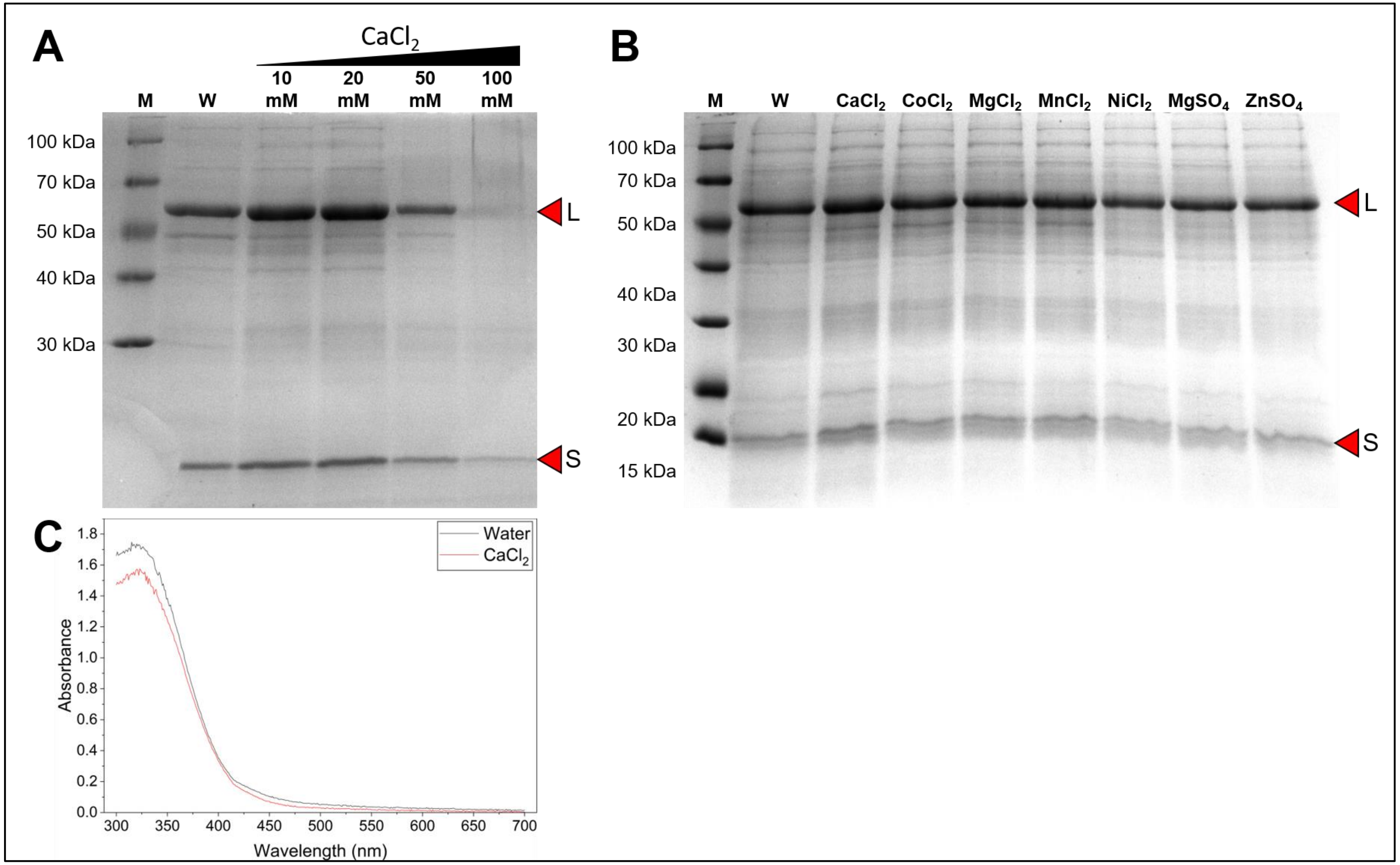
